# DeCalciOn: A hardware system for real-time decoding of *in-vivo* calcium imaging data

**DOI:** 10.1101/2022.01.31.478424

**Authors:** Zhe Chen, Garrett J. Blair, Changliang Guo, Jim Zhou, Alicia Izquierdo, Peyman Golshani, Jason Cong, Daniel Aharoni, Hugh T. Blair

**Affiliations:** Department of Electrical and Computer Engineering, UCLA, Los Angeles, CA 90095-1563, USA; Department of Psychology, UCLA, Los Angeles, CA 90095-1563, USA; David Geffen School of Medicine, University of California Los Angeles, Los Angeles, CA, 90095 USA; Department of Neurology, David Geffen School of Medicine, University of California, Los Angeles, Los Angeles, CA, USA; Integrative Center for Learning and Memory, University of California, Los Angeles, Los Angeles, CA, USA

## Abstract

Epifluorescence miniature microscopes (“miniscopes”) are widely used for *in vivo* calcium imaging of neural population activity. Imaging data is usually collected while subjects are engaged in a task and stored for later offline analysis, but emerging techniques for online imaging offer potential for novel real-time experiments in which closed-loop interventions (such as neurostimulation or sensory feedback) are triggered at short latencies in response to neural population activity. Here we introduce *DeCalciOn*, a plug-and-play hardware device for online population decoding of *in vivo* calcium signals that can trigger closed-loop feedback at millisecond latencies, and is compatible with miniscopes that use the UCLA Data Acquisition (DAQ) interface. In performance tests, the position of rats (n=13) on a linear track was decoded in real time from hippocampal CA1 population activity by 24 linear classifiers. DeCalciOn required <2.5 ms after each end-of-frame to decode up to 1,024 calcium traces and trigger TTL control outputs. Decoding was most efficient using a ‘contour-free’ method of extracting traces from ROIs that were unaligned with neurons in the image, but ‘contour-based’ extraction from neuronal ROIs is also supported. DeCalciOn is an easy-to-use system for real-time decoding of calcium fluorescence that enables closed-loop feedback experiments in behaving animals.

## Introduction

Miniature epifluorescence microscopes (“miniscopes”) can be worn on the head of an unrestrained animal to perform *in vivo* calcium imaging of neural population activity during free behavior^1,2,3,4,5^. Imaging data is usually collected while subjects are engaged in a task and stored for later offline analysis. Popular offline analysis packages such as CaImAn^20^ and MIN1PIPE^21^ employ algorithms^29^ for demixing crossover fluorescence between multiple sources to extract calcium traces from single neurons, but these algorithms cannot be implemented in real time because they rely on acausal computations. Emerging techniques for online trace extraction^6,7,8,9,10,11,26^ offer potential for carrying out real-time imaging experiments in which closed-loop neurostimulation or sensory feedback are triggered at short latencies in response to neural activity decoded from calcium fluorescence^12,14,25^. Such experiments could open new avenues for investigating the neural basis of behavior, developing brain-machine interface devices, and preclinical testing of neurofeedback-based therapies for neurological disorders^13^. To advance these novel lines of research, it is necessary to develop and disseminate accessible tools for online calcium imaging and neural population decoding.

Here we introduce *DeCalciOn*, a plug-and-play hardware device for *De*coding *Calci*um Images *On*line that is compatible with existing miniscope devices which utilize the UCLA Miniscope DAQ interface board^3,15,16,17^. Device performance was evaluated by implementing 24 linear classifiers to decode hippocampal population activity in unrestrained rats (n=13) running on a linear track. We show that the system requires <2.5 ms after each end-of-frame to decode up to 1,024 calcium traces and trigger TTL outputs that can control external devices. DeCalciOn performs real-time trace extraction by summing fluorescence in regions of interest (ROIs) assigned to each individual trace, without any demixing of crossover fluorescence. In our performance tests, decoding accuracy achieved with this simple trace extraction method matched or exceeded that obtained with offline trace extraction using constrained non-negative matrix factorization^20,29^ (CNMF). Online decoding was most accurate and efficient when traces were extracted using “contour-free” ROIs that tiled the entire image frame, and did not overlap with individual neurons. Hence, real-time decoding of neural population activity from calcium fluorescence was most efficient when minimal bandwidth was devoted to computations for source extraction that failed to improve decoding accuracy.

In summary, DeCalciOn provides the research community with a low-cost, open-source, easy-to-use hardware platform for real-time decoding of neural population activity that will permit researchers to perform novel closed-loop experiments in behaving animals. All hardware, software, and firmware are openly available through miniscope.org.

## Results

The DeCalciOn hardware system (Fig. 1) is implemented on an Avnet™ Ultra96 development board featuring a Xilinx™ Zynq Ultrascale+ multiprocessor system-on-a-chip (MPSoC) with 2GB DRAM. A custom interface board mated to the Ultra96 receives real-time image data from a modified version of the UCLA Miniscope Data Acquisition (DAQ) interface, which has 14 flywires soldered to the PCB for transmitting deserialized video to the Ultra96 (pre-modified DAQ boards can be obtained through miniscope.org). Results presented here were obtained using a Large Field-of-View (MiniLFOV) version^17^ of the UCLA Miniscope featuring 1296×972 pixel resolution acquisition at 22.8 frames-per-second (fps).

**Figure 1.**
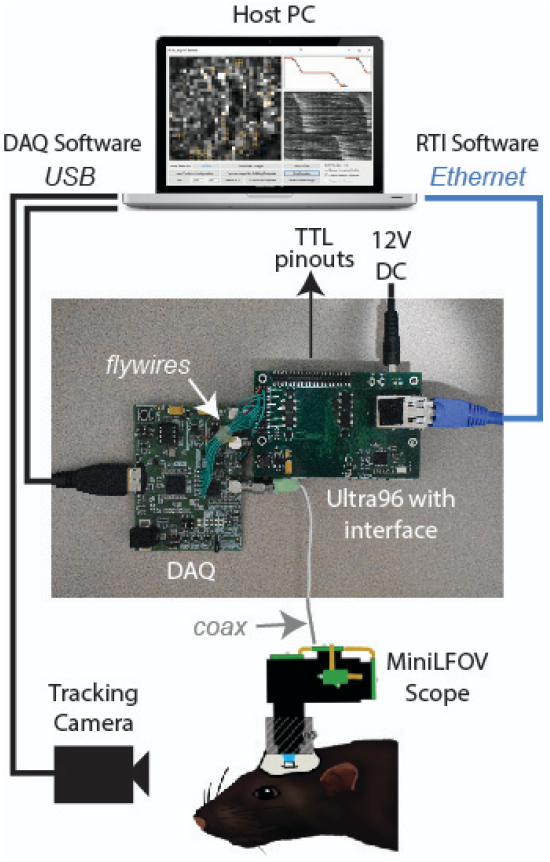
System Hardware. Miniscope connects to DAQ via coax cable, DAQ connects to Ultra96 via flywires, host PC connects to Ultra96 via Ethernet and to DAQ via USB 3.0; TTL pinouts from Ultra96 can drive closed-loop feedback stimulation from external devices.

### Image processing pipeline

Incoming frames from the MiniLFOV were cropped to a 512×512 subregion containing the richest area of fluorescing neurons in each rat, and stored to BRAM for real-time processing on the Ultra96 (Fig. 2, bottom left). Online processing of each image frame was performed in four sequential steps: 1) motion stabilization, 2) background removal, 3) calcium trace extraction, and 4) decoding. Steps 1-3 were performed by our custom ACTEV (Accelerator for Calcium Trace Extraction from Video) firmware running in the programmable logic fabric of the MPSoC’s field-programmable gate array (FPGA). Step 4 was performed by a C# program running under the FreeRTOS operating system on the MPSoC’s embedded ARM core.

**Figure. 2.**
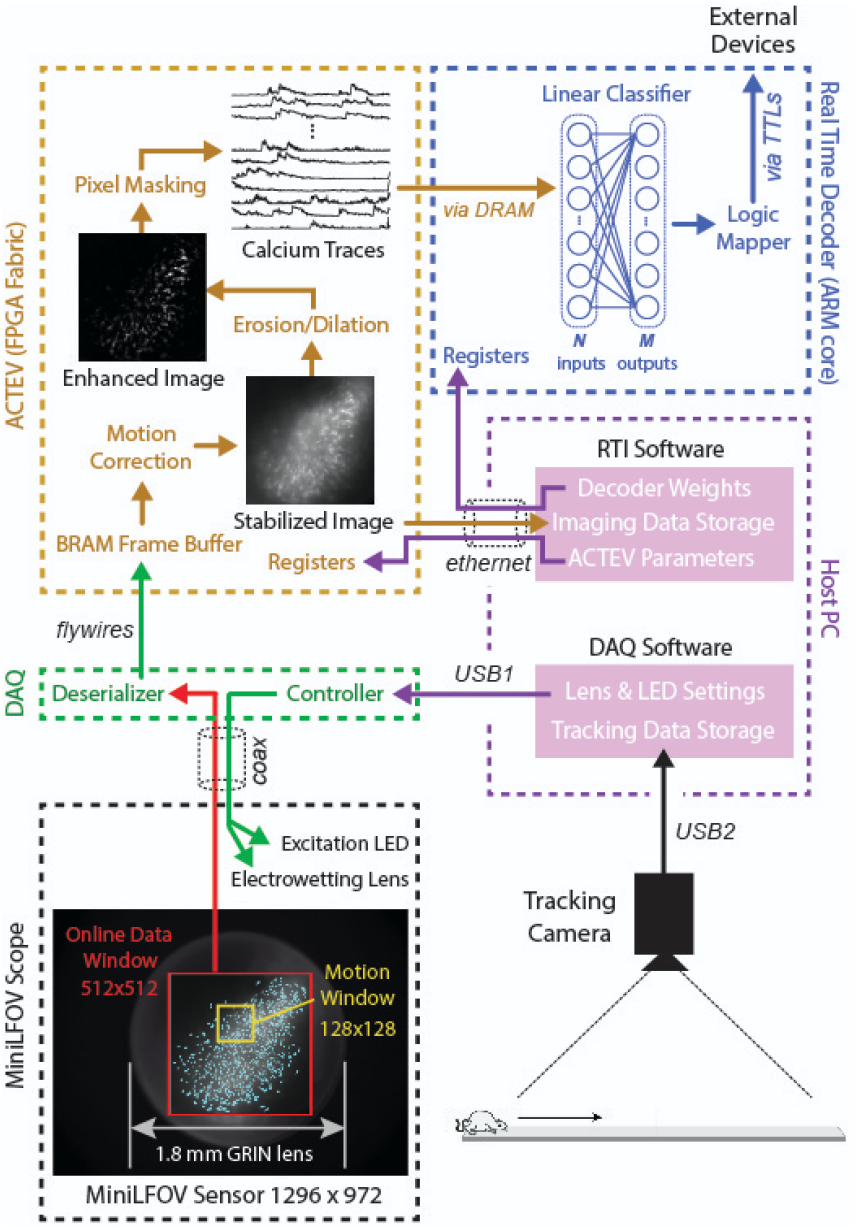
Online imaging and control pipelines. Serialized video data from the MiniLFOV is transmitted through a coaxial tether cable to the DAQ, where it is deserialized and transmitted via the flywire bus to ACTEV (Accelerator for Calcium Trace Extraction from Video) firmware programmed on the FPGA of the Ultra96. When used with MiniLFOV (as in performance tests reported here), ACTEV crops the incoming 1296×972 sensor image down to a 512×512 sub-window (selected to contain the richest area of fluorescing neurons in each rat) before storing video frames to a BRAM buffer on the FPGA. Subsequent motion correction and calcium trace extraction steps are described in the main text. Extracted traces are stored to DRAM from which they can be read as inputs to a linear classifier decoder running on the Ultra96 ARM core. Decoder output is fed to a logic mapper that can trigger TTL output signals from the Ultra96, which in turn can control external devices such as stimulators for generating closed-loop feedback.

To correct for translational movement of brain tissue (Step 1), a 128×128 pixel area with distinct anatomical features was selected within the 512×512 cropped subregion to serve as a motion stabilization window. ACTEV’s image stabilization algorithm^8^ rigidly corrects for translational movement of brain tissue by convolving the 128×128 stabilization window in each frame with a 17×17 contrast filter kernel, and then applying a fast 2D FFT/IFFT based algorithm to correlate the window contents with a stored reference template (derived at the beginning of each experimental session) to obtain a 2D motion displacement vector for the current frame. Supplementary Video 1 demonstrates online performance of ACTEV’s real-time motion stabilization algorithm.

After motion stabilization, ACTEV removes background fluorescence (Step 2) from the 512×512 image by performing a sequence of three enhancement operations^9^: smoothing via convolution with a 3×3 mean filtering kernel, estimating the frame background via erosion and dilation with a 19×19 structuring element^22^, and subtracting the estimated background from the smoothed image. These operations produce an enhanced image in which fluorescing neurons stand out in contrast against the background (see Supplementary Video 1; “Enhanced Image” in Fig. 2). The enhanced image then is filtered through a library of up to 1,024 binary pixel masks (each up to 25×25 in size) that define ROIs within fluorescence is summed to extract calcium traces (Step 3); each mask can be centered anywhere in the 512×512 window. Pixel masks can be created using either a *contour-based* or a *contour-free* approach (see below).

Population vectors of extracted calcium trace values are stored to the DRAM, and then decoded (Step 4) by sending them as inputs to a linear classifier running on the MPSoC’s ARM core. The linear classifier’s output vector during each image frame is relayed to a logic mapper which can trigger voltage changes on up to 8 TTL pinouts from the Ultra96, and thereby control closed-loop neural or sensory feedback from external devices in response to decoded calcium fluorescence signals. Throughout online imaging sessions, the DeCalciOn system is controlled from a host PC running standard Miniscope DAQ software (to focus and adjust the MiniLFOV) alongside newly developed real-time interface (RTI) software to communicate with ACTEV (see below). Raw video data (including real-time motion correction vectors) and calcium trace values are transmitted via ethernet from the Ultra96 to a host PC for storage, so that additional analyses of calcium data can later be performed offline.

### Real-time position decoding from CA1 place cells

Device performance was evaluated using image data collected from the hippocampal CA1 region while Long-Evans rats (n=13) ran back and forth on a 250 cm linear track (Fig. 3A). CA1 pyramidal neurons behave as “place cells” that fire selectively when an animal traverses specific locations in space^18^, so a rodent’s position can be reliably decoded from CA1 population activity^2,3,19,28^. For performance testing, we used a virtual sensor (see Methods) that was capable of feeding stored image data to the ACTEV firmware at 22.8 fps, exactly as if raw video data were arriving from the MiniLFOV in real time. This allowed different firmware algorithms (for example, contour-based versus contour-free decoding; see below) to be compared and benchmarked on the same stored datasets. Results obtained with the virtual sensor were verified to be identical with those obtained in real-time.

**Figure 3.**
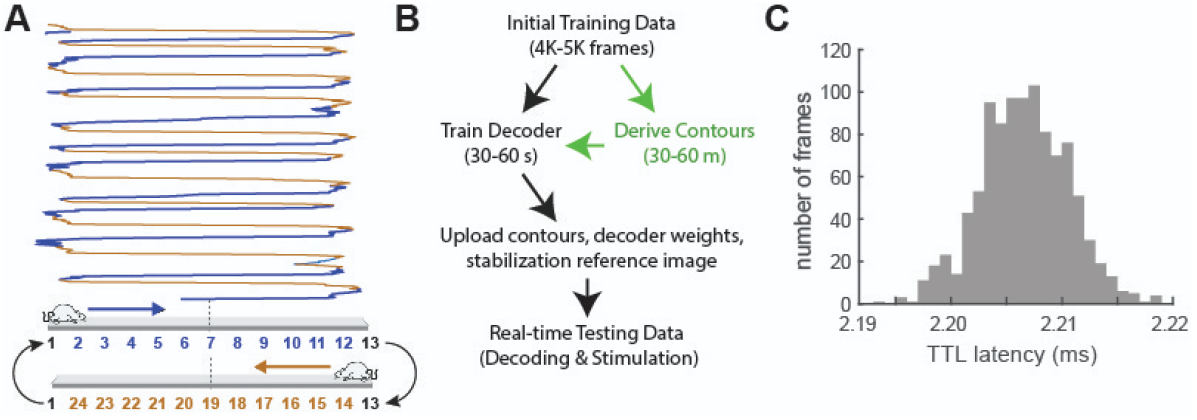
Real-time position decoding. *A*) The CA1 layer of the hippocampus was imaged while rats ran laps on a 250 cm linear track, subdivided into 24 position bins. *B*) Sequence of experimental steps for a real-time imaging & decoding session. *C)* Latency distribution for decoding 1,024 online calcium traces.

During a typical real-time session, the experimenter carries out four steps (Fig. 3B): 1) collect an initial imaging dataset and store it to the host PC, 2) pause for an intermission to identify cell contours if necessary (this is only required for contour-based decoding; see below) and train a linear classifier to decode behavior from the initial dataset, 3) upload classifier weights from the host PC to the Ultra96, and 4) perform real-time decoding with the trained classifier. To mimic these steps using the virtual sensor in our performance tests, one session of image data was collected and stored from each of the 13 rats, yielding ∼7 min (8K-9K frames) of sensor and position tracking data per rat. The linear classifier was then trained on data from the first half of each session, and tested on data from the second half.

When place cell activity is analyzed or decoded offline (rather than online) from previously stored neural and position data, it is common practice to perform *speed filtering* that omits time periods when the rat is sitting still. This is done because during stillness, the hippocampus enters a characteristic “sharp wave” EEG state during which place cell activity is less reliably tuned for the animal’s current location^24^. Here, speed filtering was not implemented during online decoding of CA1 fluorescence because the linear classifier’s input layer received only real-time calcium trace data, and not position tracking data (which would be needed for speed filtering). It is possible that some speed filtering may have been implicitly learned by the decoder if information about the animal’s speed was encoded in any of the calcium traces, but explicit speed filtering of calcium data was not performed prior to training or testing the online classifier. Despite this, DeCalciOn was able to achieve highly accurate decoding of the rat’s position from calcium traces (see below).

After the decoder had been trained, real-time classifier output from each frame was used to trigger TTL feedback outputs from the Ultra96. The latency to generate feedback from decoded calcium traces was measured as the time elapsed between acquiring the last pixel of each cropped frame and the rising edge of the triggered TTL pulse. Under a worst-case scenario of decoding the maximum number of calcium traces supported by the hardware (N=1,024), mean decoding latency was 2,206±4 *μ*s and never exceeded 2,220 *μ*s (Fig. 3C). This highlights one of the main advantages inherent in our FPGA-based hardware design; it would be difficult to trigger feedback with such short latencies and low variability if online image data were processed on a CPU or GPU running programs with multiple threads, or if real-time video were relayed to the image processor through USB or ethernet instead of by a direct hardware connection with the DAQ.

### Contour-based versus contour-free decoding

Decoding performance was compared for two different methods of extracting online fluorescence traces: *contour-based* versus *contour-free*. Contour-based extraction derived calcium traces by summing fluorescence within pixel masks containing cell contours (Fig. 4A; Supplementary Video 2) identified using constrained non-negative matrix factorization (CNMF) by the CaImAn pipeline^20^, which was applied to the training dataset during the intermission period. After contours had been identified in this way, the training dataset was fed through a simulator to reconstruct online calcium traces that would have been extracted in real time during the first half of the session using the contour masks; simulated traces were then used to train the decoder. In contrast with contour-based extraction, contour-free extraction summed fluorescence within pixel masks obtained by partitioning the 512×512 image into a 32×32 sheet of tiles, excluding 124 tiles bordering the frame to prevent artifacts from motion stabilization (Fig. 4B; Supplementary Video 3). The contour-free method always yielded a total of 900 calcium traces per rat, whereas the contour-based method yielded a variable number of traces depending upon the number of neurons detected by CaImAn.

**Figure 4.**
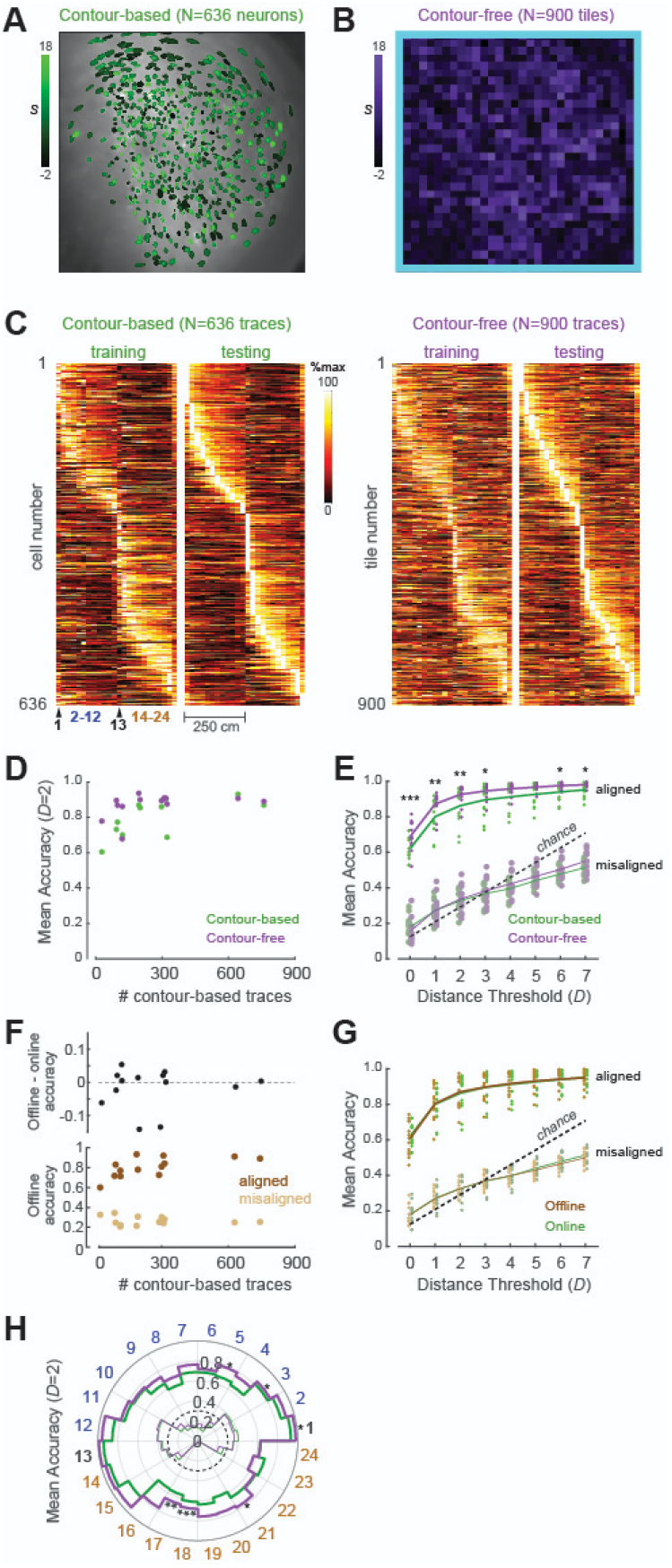
Analysis of decoding accuracy. *A*) Cell contours identified by CAIMAN in one rat; intensity of green shading shows the similarity score, *S*, for traces exracted from each contour. *B*) Mosaic of square pixel mask tiles used for contour-free trace extraction in the same rat as ‘A’; Intensity of purple shading indicates *S* value for traces extracted from each tile. Edge tiles (light blue border) were omitted from trace extraction. *C*) Tuning curve heatmaps from training and testing epochs for contour-based (left) and contour-free (right) traces extracted from the rat shown in ‘A’ and ‘B’; rows are sorted by location of peak activity in the testing epoch. *D*) Scatter plot shows accuracy of decoding from contour-free or contour-based traces (with *D*=2) as a function of the number of contour-based traces in each rat. *E*) Mean decoding accuracy (y-axis) over all position bins as function of *D* (x-axis) for each rat (dots) and averaged over rats (lines), contour-free and contour-based classifers were both significantly more accurate than chance or control classifiers trained on calcium traces misaligned with position data (p<.0001 for all *D*). *F*) *Top*. Scatter plot shows offline minus online accuracy for contour-based traces; positive y-values indicate more accurate decoding from offline traces. *Bottom:* Offline contour-based decoding accuracy for classifier trained on traces that were aligned or misaligned (control) with position data *G*) Mean contour-based decoding accuracy was similar for offline and online traces derived from the same contours (p>.45 for all *D*). *H*) Polar plot of mean decoding accuracy (radius) per position bin (angle) for *D*=2. Asterisks in ‘E’ and ‘H* show significance levels for comparison between contour-based vs. contour-free decoding. *p<.05, **p<.01. ***p<.001.

After calcium traces had been extracted from the training dataset by either method, they were aligned with position tracking data and used to train the linear classifier during the intermission period. ACTEV can utilize a real-time spike inference engine to convert raw calcium trace values into inferred spike counts prior to decoding. However, for both contour-based and contour free trace extraction methods, the linear classifier was more accurate at learning to decode the rat’s position from raw calcium traces than from inferred spikes (Supplementary Fig. 1). Similar results have been reported in prior studies of position decoding from place cell calcium fluorescence^27^. When information is decoded from raw calcium traces, the GCamp molecule effectively performs temporal integration of neural activity at a time constant similar to its decay rate. Hence, the finding that decoding is more accurate from raw traces than inferred spike counts can be interpreted to mean that when decoding position from place cell calcium signals, the ideal time constant over which to integrate neural activity is closer to the decay constant of the calcium indicator (several hundred ms for the GCamp7s indicator used here^23^) than to the video frame acquisition interval (∼44 ms for the MiniLFOV scope used here).

Training the classifier on 4K-5K frames of calcium trace data required <60 s of computing time on the host PC during the intermission period. However, contour-based trace extraction required an additional 30–60 min to identify cell contours and simulate online traces before training the classifier. Faster methods for contour identification may be possible, but contour-free trace extraction has the advantage of eliminating this delay altogether. As shown below, this advantage came at no cost to decoding accuracy in our performance tests.

After the linear classifier had been trained, learned weights were uploaded from the host PC to the Ultra96; image data from the second half of the experimental session was then fed through the virtual sensor for real-time decoding. The classifier’s output vector consisted of 23 units that used an ordinal scheme (see Methods) to represent 24 binned locations (12 per running direction) along the track (Fig. 3A, bottom). Accurate decoding requires that calcium traces must retain stable spatial tuning across the training and testing epochs of the session; to verify that this was the case, spatial tuning curves were derived for calcium traces during the first (training) versus second (testing) half of each session (Fig. 4C). A similarity score (*S*) was computed for each trace (Fig. 4A,B) using the formula *S* = -log_10_ (*P*)×sign(*R*), where *R* is the correlation between the trace’s tuning curves from training versus testing, and *P* is the significance level for *R*. Contour-based traces had higher mean *S* scores than contour-free traces in every rat (paired *t*_12_=7.02, p=1.4e^-5^); hence, at the level of individual traces, spatial tuning was better for contour-based than contour-free traces (Supplementary Fig. 2). However, the number of contour-free traces (900 per rat) was always greater than the number of contour-based traces (which varied by rat); hence, at the population level, contour-free traces could often convey more information about position than contour-based traces. To compare decoding accuracy between contour-based and contour-free traces, we measured the percentage of frames from the testing epoch during which the trained decoder’s position prediction was within ±*D* bins of the true position. Averaged over rats, the mean decoding accuracy was significantly greater for contour-free than contour-based decoding at most values of *D* (Fig. 4E,H). Analogous results have been reported in electrophysiology, where decoding is sometimes more accurate from “cluster-free” multiunit activity than from single-unit spikes^21^. Decoding position from CaImAn’s offline (demixed) traces did not improve accuracy over decoding from online traces derived from the same cell contours (Fig, 4F,G). This demonstrates that the inferior accuracy of contour-based decoding was not rooted in any loss of fidelity incurred by the transition from offline (demixed) to online (non-demixed) trace extraction.

Although mean decoding accuracy was higher for contour-free than contour-based traces in almost every rat, the contour-free advantage was greatest in rats with <400 contour-based traces (Fig. 4D), indicating that the primary reason why contour-free traces provided better decoding accuracy was because they were greater in number. CaImAn’s sensitivity parameters can be adjusted to detect more contour-based traces in each rat, and when this was done, the accuracy of contour-free and contour-based decoding became much more similar (Supplementary Fig. 3). Hence, contour-free traces provided better decoding accuracy than contour-based traces only when they were greater in number. Based on these results, the contour-free approach is recommendable over the contour-based approach as a faster and more efficient method for online trace extraction, since it is just as accurate and obviates the need to identify cell contours before training the decoder.

## Discussion

Here we have introduced DeCalciOn, an easy to use open-source hardware platform for online calcium imaging that can perform real-time decoding of neural population activity. The system is capable of triggering TTL outputs to generate neural or sensory feedback at millisecond latencies, and is therefore appropriate for use in a wide range of closed-loop neuroscience applications.

### Comparison with other platforms

Several online calcium fluorescence trace extraction algorithms have been proposed, but some of these have been validated only by offline experiments (showing that they can generate online traces that rival the quality of offline traces) without being performance tested on specific hardware platforms during real-time imaging or decoding experiments^6,11,26^.

Liu et al.^25^ introduced a system for real-time decoding with the UCLA miniscope that, unlike DeCalciOn, requires no additional hardware components; the system is implemented entirely by software running on the miniscope’s host PC. It was shown that this system can implement a single binary SVM classifier to decode calcium traces from 10 regions of interest (ROIs) in mouse cortex at latencies <3 ms. However, the system does not perform online motion stabilization, which could degrade performance when large numbers of ROIs are spaced closely together. Any system that lacks motion stabilization would also be vulnerable to artifactually decoding behavior from brain motion (which can be correlated with behavior) rather than neural activity. Liu et al.^25^ did not report how their system’s decoding latency scaled with the number of classifiers or the size/number of ROIs, but if serial processing on the host PC scales linearly with these variables, then decoding latencies of several seconds or more would be incurred when implementing 24 classifiers to decode 1,204 calcium traces.

Zhang et al.^14^ performed 2-photon calcium imaging experiments that incorporated real time image processing (running on a GPU) to detect neural activity and trigger closed-loop optical feedback stimulation in mouse cortex. With online motion correction enabled, the system achieved mean feedback latencies of 8.5+*n*/30 ms after the end of each frame, where *n* is the number of ROIs in the image. This system would thus be expected to incur a mean decoding latency of ∼43 ms to extract traces from 1,204 ROIs; the system’s variability appears to scale with latency, so delays might be >60 ms for some frames and <20 ms for others. Zhang et al.’s^14^ system is well suited for experiments in which fast (<10 ms) feedback is triggered from tens of ROIs or slower feedback (40±20 ms) is triggered from hundreds of ROIs. By contrast, DeCalciOn can achieve feedback latencies of <2.5 ms with submillisecond variability even for large ROI counts (Fig. 3C), and would thus be preferable for experiments requiring fast closed-loop feedback triggered from large numbers of ROIs.

### Contour-free source extraction

In our experiments, contour-free outperformed contour-based decoding when the number of contour-based traces was less than half the number of contour-free traces (Fig. 4D). When the number of contour-based and contour-free traces was similar, both methods yielded similar decoding accuracies (Supplementary Fig. 3). Decoding from offline (demixed) contour-based traces was not more accurate than decoding from online (non-demixed) contour-based traces (Supplementary Fig. 2), so there was no evidence that decoding accuracy suffered at all from greater crosstalk between sources during online trace extraction. In the process of ‘demixing’ fluorescence signals from one another, CaImAn’s CNMF algorithm does not in any way consider whether pixels contain information about behavior, so it may be prone to discard pixels that contain decodable position information, especially when using conservative parameters for contour detection. Supporting this, contour-free pixel masks with high *S* scores (bright purple squares in Fig. 4B) were sometimes located in regions of the image where CaImAn detected no cell contours at all under conservative sensitivity parameters (green pixel masks in Fig. 4A). Based on these results, the contour-free approach is recommendable as the efficient online trace extraction method for real-time population decoding with DeCalciOn.

### Summary and Conclusions

DeCalciOn’s low cost, ease of use, and latency performance compare favorably against other real time imaging systems proposed in the literature. By making DeCalciOn widely available to the research community, we provide a platform for real-time decoding of neural population activity that we hope will facilitate novel closed-loop experiments and accelerate discovery in neuroscience and neuroengineering. All of DeCalciOn’s hardware, software, and firmware are openly available through miniscope.org.

## Acknowledgements

The authors thank Xilinx Corporation for donation of the Ultra96 board. We also thank Jeffrey Johnson for technical assistance in designing the custom interface board, and Shiyun Wang and Ryan Grgurich for helpful comments on the manuscript. This work was supported by NSF NeuroNex 17040708 (HTB, PG, DA, JC), 1UF1NS107668 (DA), and MH122800 (AI, HTB).

## Materials and Methods

### Hardware

The DeCalciOn system is designed for use with UCLA Miniscope devices^3^ and other miniscopes^15,16,17^ that use the UCLA DAQ interface. The DAQ hardware requires modification by soldering flywires to the PCB (see Fig. 1); instructions for doing this are available at https://github.com/zhe-ch/ACTEV and pre-modified DAQ boards are available at miniscope.org. DeCalciOn is implemented on an Avnet Ultra96 development board (available from the manufacturer at Avnet.com), which must be mated to our custom interface board available at miniscope.org (the interface can also be ordered from a PCB manufacturer with design files available at https://github.com/zhe-ch/ACTEV). ACTEV firmware (for online image stabilization, image enhancement, calcium trace extraction) and embedded software (for real-time decoding and ethernet communication, which runs under the FreeRTOS operating system) can be programmed onto the Ultra96 board by copying a bootable bitstream file to a MicroSD card and then powering up the board with the card inserted. The MiniLFOV version of the bitstream file (used for experiments reported here) is available at https://github.com/zhe-ch/ACTEV. Bitstream files for other miniscope models, as well as Vivado™ HLS C source code for generating new custom bitstream files, are available on request.

### Software

Real-time imaging and decoding are controlled by custom RTI software (available at https://github.com/zhe-ch/ACTEV) which runs on the host PC alongside standard DAQ software (available at miniscope.org) for adjusting the gain and focus of the Miniscope device. The DAQ software’s default operation mode is to receive and display raw Miniscope video data from the DAQ via a USB 3.0 port (and store this data if the storage option has been selected), and also to receive and display raw behavior tracking video from a webcam through a separate USB port. At the start of a real experimental session, data acquisition by both programs (DAQ and RTI software) is initiated simultaneously with a single button click in the RTI user interface, so that Miniscope video storage by the RTI software and behavior video storage by the Miniscope DAQ software are synchronized to begin at exactly the same time. This allows behavioral data stored by the DAQ software to be aligned with Miniscope video and calcium trace data stored by the RTI software. During the intermission period between initial data acquisition and real-time inference, data stored by DAQ and RTI software is used to train the linear classifier on the host PC. Trained classifier weights are then uploaded to the Ultra96 for real-time decoding.

### Contour-based pixel masks

Pixel masks for contour-based trace extraction were derived using the CaImAn^20^ pipeline (implemented in python) to analyze motion-corrected sensor images from the training dataset (as noted in the main text, this took 30-60 min of computing time on the host PC). CaImAn is an offline algorithm that uses constrained non-negative matrix factorization (CNMF) to isolate single neurons by demixing their calcium traces. The CNMF method is acausal so it cannot be used to extract traces in real time. But during offline trace extraction, CaImAn generates a set of spatial contours identifying pixel regions from which each demixed trace was derived, which ACTEV then uses as pixel masks for online extraction of contour-based traces. Once pixel masks were identified from the training data, motion-correct miniscope video from the training period was passed through an offline simulator that used ACTEV’s causal algorithm for extracting calcium traces from contour pixel masks. This yielded a set of calcium traces identical to those that would have been extracted from the training data in real time by ACTEV. These simulated traces were then used as input vectors to train the linear classifier. Since the online traces are not generated by CNMF (and are thus not demixed from one another), they are susceptible to contamination from fluorescence originating outside of contour boundaries. However, this crosstalk fluorescence did not impair decoding since similar accuracy results were obtained by training the linear classifier on online or offline contour-based traces (Fig. 4F,G).

### Contour-free pixel masks

To implement contour-free trace extraction, we simply partitioned the 512×512 image frame into a 32×32 sheet of tiles, each measuring a square of 16×16 pixels (Fig. 4B). No traces were extracted from 124 tiles bordering the edge of the frame, to avoid noise artifacts that might arise from edge effects in the motion stabilization algorithm. Hence, a total of 1,024-124=900 pixel mask tiles were used for contour-free calcium trace extraction. These traces were derived in real time and stored to the host PC throughout the initial data acquisition period, so they were immediately available for training the linear classifier at the start of the intermission. Consequently, an advantage of contour-free trace extraction is that the intermission period between training and testing is shortened to just a few minutes, because the lengthy process of contour identification is no longer required. A disadvantage of contour-free trace extraction is that contour tiles do not align with individual neurons in the sensor image. As reported in the main text, this lack of alignment between neurons and pixel masks did not impair (and often enhanced) position decoding; however, contour-free decoding does place limits upon what can be inferred about how single neurons represent information in imaged brain regions.

### Fluorescence summation

After pixel masks were created using one of the two methods (contour-based or contour-free) and uploaded to the FPGA, calculations for extracting calcium traces from the masks were the same. Each mask specified a set of pixels over which grayscale intensities were summed to obtain the fluorescence value of a single calcium trace:

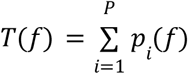

where *T(f)* is the summed trace intensity for frame *f*, and *p*_*i*_(*f*) is the intensity of the *i*^th^ pixel in the mask for frame *f*, and *P* is the number of pixels in the mask. Each contour-free mask was a square tile containing 16×16=256 pixels (Fig. 4B), whereas the size of each contour-based mask depended upon where CaImAn identified neurons in the image (Fig, 4A).

### Drop filtering

The mean size of contour-based masks was 20-50 pixels (depending on the rat), which was an order of magnitude smaller than contour-free tiles. In a few rats, a small amount of jitter sometimes penetrated the online motion correction filter, causing the stabilized sensor image to slip by 1-2 pixels against stationary contour masks during 1-2 frames (see Supplementary Video 1, line graph at lower right). This slippage misaligned pixels at the edges of each contour mask, producing intermittent noise in the calcium trace that was proportional to the fraction of misaligned pixels in the contour, which in turn was inversely proportional to the size of the contour, so that motion jitter caused more noise in traces derived from small contour masks than large contour masks. To filter out this occasional motion jitter noise from calcium traces, a *drop filter* was applied to traces derived from contour masks that contained fewer than 50 pixels. The drop filter exploited the fact that genetic calcium indicators have slow decay times, and therefore, sudden drops in trace fluorescence can be reliably attributed to jitter noise. For example, the GCamp7s indicator used here has a half decay time^18^ of about 0.7 s, so at the MiniLFOV’s 22.8 Hz frame rate, a fluorescence reduction of more than 5% between frames can only arise from jitter artifact. The drop filter defines a maximum permissible reduction in fluorescence between successive frames as follows:

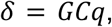

where *G* is the maximum possible fluorescence intensity for any single pixel (255 for 8-bit grayscale depth), *C* is the number of pixels in the contour mask for the trace, and *q* is a user-specified sensitivity threshold, which was set to 0.9 for all results presented here. The drop-filtered calcium trace value for frame *f* was given by:

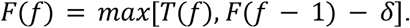

It should be noted that while drop filtering protects against artifactual decreases in trace values, it offers no protection against artifactual increases that might masquerade as neural activity events. However, motion jitter almost never produced artifactual increases in fluorescence for the small contours to which drop filtering was applied, because small contours were “difficult targets” for patches of stray fluorescence to wander into during jitter events that rarely exceeded 1-2 pixels of slippage. Larger contours were slightly more likely to experience artifactual fluorescence increases during jitter events, but in such cases, the artifact was diluted down to an inconsequential size because it only affected a tiny fraction of the large contour’s pixels. In summary, jitter artifact was highly asymmetric, producing artifactual decreases but not increases for small contours, and producing negligible artifact of any kind for large contours.

### Spike inference

Once trace values have been extracted (and drop filtered if necessary), ACTEV can apply a real-time spike inference engine to convert raw calcium trace values into inferred spike counts for each frame. Briefly, the spike inference engine measures how much the trace value in the current frame has changed from the prior frame, *T*(*f*)-*T*(*f-1*), and compares this against a threshold value, Φ, which is equal to 2.5 times the standard deviation of difference values between successive frames of the same trace in the training dataset. For the CA1 data reported here, we found that the linear classifier was more accurate at decoding raw (unconverted) calcium traces than inferred spike counts (Supplementary Fig. 1). Decoding from raw calcium traces is not only more accurate but also more computationally efficient; therefore, the main text reports results obtained by decoding raw calcium trace values.

**Supplementary Fig. 1.**
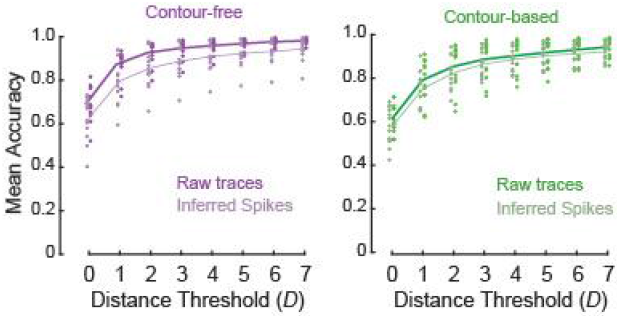
Decoding from raw calcium traces versus inferred spikes. For contour-free (left) and contour-based (right) trace extraction methods, accuracy of position decoding was higher when the linear classifier was trained and tested on raw calcium trace values than inferred spike events.

### Training the linear classifier

The linear classifier has *N* inputs and *M* outputs, where *N* is the number of calcium traces. In performance tests reported here, the output layer utilized an ordinal scheme to represent each of the 24 binned track locations: at position bin #1 the target output for all output units was +1, at position bin #2 the target output was -1 for the first output unit and +1 for the remaining units, at position bin #3 the target output was -1 for the first two output units and +1 for the remaining units, and so on. Under this ordinal representation scheme, the total number of output units is equal to *M*=*K*-1, where *K* is the number of categories (position bins) to be encoded. Since the linear track was subdivided into *K*=24 spatial bins, there were *M*=24-1=23 binary classifier units in the decoder’s output layer. To train the linear classifier, data stored during the initial acquisition period by the DAQ and RTI software was analyzed during the intermission period by custom MATLAB scripts, available at https://github.com/zhe-ch/ACTEV. Training data consisted of calcium trace input vectors (derived by either contour-based or contour-free methods described above) and target position output vectors (encoding the rat’s true position) for each Miniscope frame. These training vectors were passed to MATLAB’s *fitceoc* function (from the machine learning toolbox) to compute the trained linear classifier weights, which were then uploaded from the host PC to the Ultra96 via ethernet, so that the real-time classifier could subsequently decode calcium traces in real time. As a control condition, calcium traces and tracking data were circularly shifted against one another by 500, 1000, 1500, 2000, or 2500 frames before training, and the trained classifier was then tested on correctly aligned trace inputs and target position outputs. Accuracy results from all five shift values were averaged together to obtain the ‘misaligned’ decoding accuracies plotted in Figs. 2E,F of the main text.

### Decoding calcium traces

Once classifier weights had been uploaded to the Ultra96 from the host PC, it was then possible for the embedded ARM core in the Ultra96 MPSoC to read in calcium trace values from the DRAM in real time as each frame of data arrived from the Miniscope, and present these values as input to the linear classifier. The linear classifier performed calculations equivalent to MATLAB’s *predict* function (from the machine learning toolbox) to generate an output vector predicting the rat’s position on the track during each frame. This prediction was sent back to the host PC via ethernet for storage, and was also sent to a programmable *logic mapper* running on the ARM core, which converted the linear classifer’s output vector into a pattern of square pulses generated at on of 8 TTL pinouts from the Ultra96. TLL outputs are capable of driving closed-loop neural or sensory feedback via external devices such as lasers, electrical stimulators, or audiovisual display equipment. For performance testing, we measured the latency to generate TTL outputs from decoded neural activity (see Fig. 1H in the main text), but we did not deliver closed loop feedback to our pilot rats. As explained in the Results, decoding from contour-free traces was more accurate than from contour-based traces. This was because there were more contour-free traces in every rat (Supplementary Fig. 2).

**Supplementary Fig. 2.**
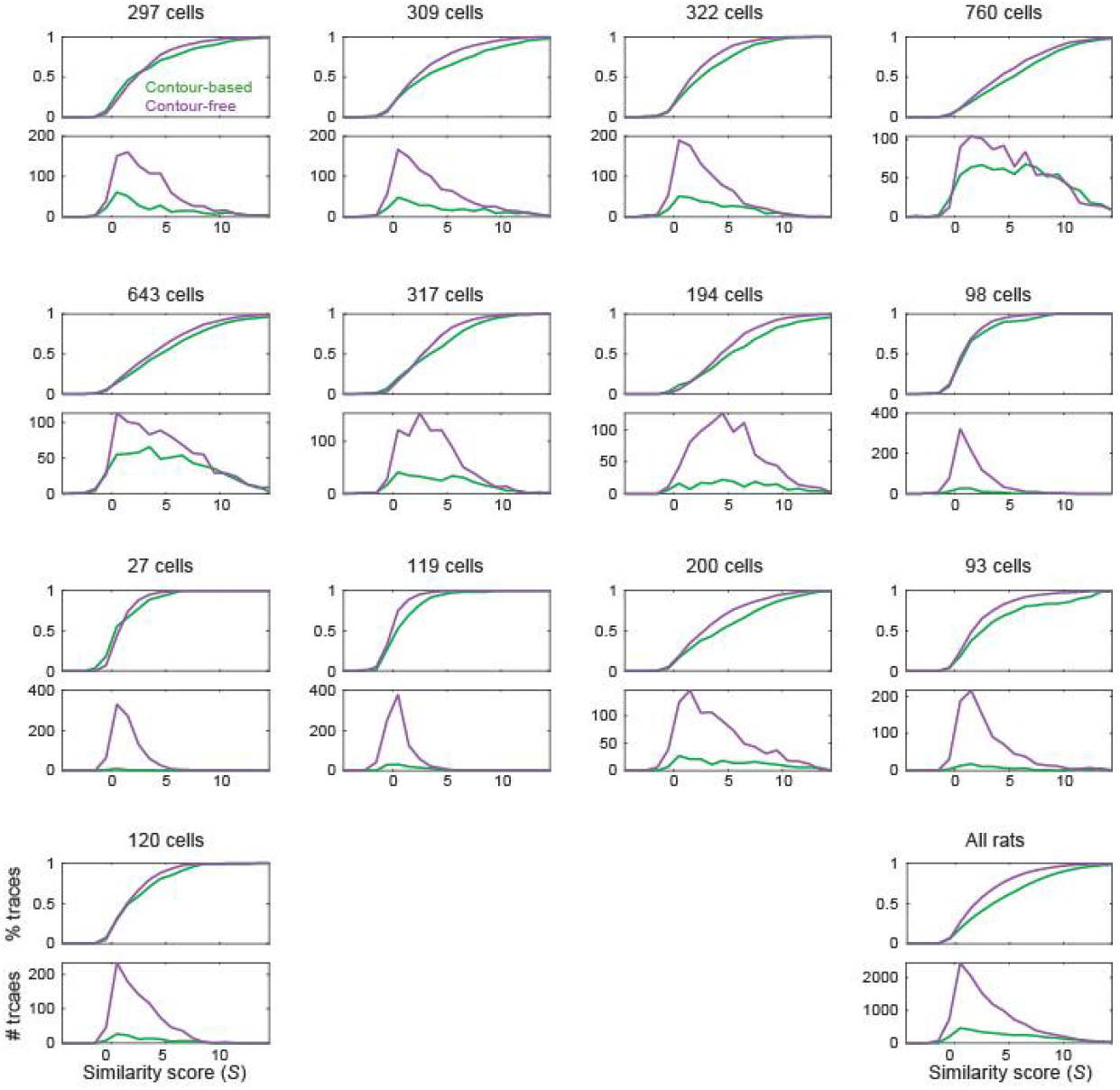
Spatial similarity score distributions by rat. Each plot shows the cumulative probability density (top) and frequency distribution (bottom) of *S* scores from one rat; lower right plot shows *S* scores pooled over all rats. In every rat, the cumulative S distribution for contour-based traces is shifted to the right, indcating that contour-based traces were generally of higher quality (had more stable spatial tuning) than contour-free traces. But the number of contour-free traces (900 per rat) was always greater than the number of contour-based traces (shown at the top of each plot), and this was equally true for traces with high and low *S* values. Therefore, for position decoding, the lesser quality of individual contour-free traces was compensated by the fact that were greater in number, which was beneficial for population decoding.

**Supplementary Fig. 3.**
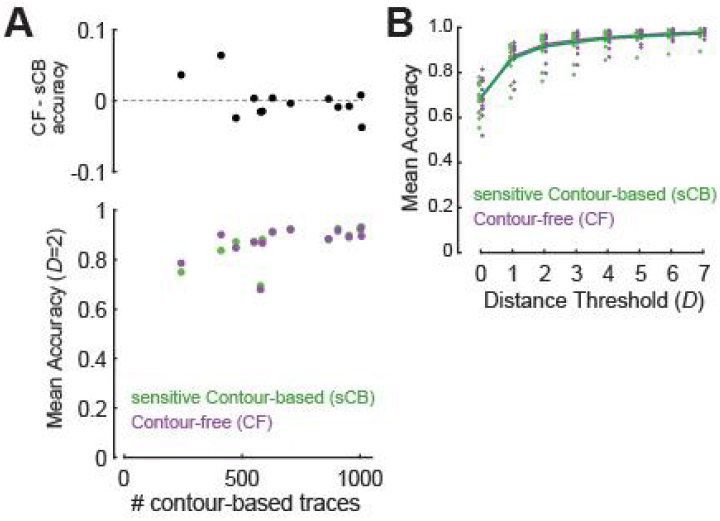
Increasing the number of contour-based traces improves decoding accuracy. *A*) CAIMAN senstivity for contour detection was increased, resulting in larger numbers of contour-based (sCB) traces being detected (x-axis). At *D*=2, decoding accuracy for sCB and contour-free (CF) traces was nearly identical in most rats, although CF decoding was still slightly more accurate in rats with the fewest sCB traces. *A*) Mean decoding accuracy for sCB and CF traces was similar for all values of *D*.

### Spatial tuning curves

To generate spatial tuning curves for fluorescence traces, each trace was first referenced to zero by subtracting it’s minimum observed fluorescence value (over all frames) from the value in every frame: *Z*(*f*) = *T*(*f*) − *min*(*T*), where *Z*(*f*) is the zero-referenced trace value for frame *f*. The 250 cm linear track was subdivided into 24 evenly spaced spatial bins (12 each in the left-to-right and right-to-left directions; see Fig. 2A). Fluorescence activity, *A*, in each bin *b* was averaged by

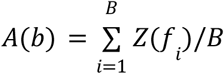

where *B* is the number of visits to bin *b* and *f*_*i*_ is the *i*^th^ frame during which the rat was visiting bin *b*. The tuning curve vector [*A*_1_, *A*_2_, …, *A*_24_] was then normalized by dividing each of its elements by the value of the maximum element. Normalized tuning curve vectors were plotted as heatmaps in Figs. 2C and 2D. Spatial similarity scores (*S*) were computed as described in the main text; *S* distributions for each rat are shown in Extended Data Fig. 3.

### Subjects

13 Long Evans rats (7F,6M) were acquired from Charles River at 3 months of age. Subjects were singly-housed within a temperature and humidity controlled vivarium on a 12 hour reverse light-cycle. Surgical procedures began after a one week acclimation period in the vivarium, and recordings and behavioral experiments began around 5 months of age. All experimental protocols were approved by the Chancellor’s Animal Research Committee of the University of California, Los Angeles, in accordance with the US National Institutes of Health (NIH) guidelines.

### Surgical procedures

Subjects were given two survival surgeries prior to behavior training in order to record fluorescent calcium activity from hippocampal CA1 cells. During the first surgery, rats were anesthetized with 5% isoflurane at 2.5 L/min of oxygen, then maintained at 2-2.5% isoflurane while a craniotomy was made above the dorsal hippocampus. Next, 1.2 *μ*L of AAV9-Syn-GCamp7s (AddGene) was injected at .12*μ*L/min just below the dorsal CA1 pyramidal layer (−3.6 AP, 2.5 ML, 2.6 DV) via a 10uL Nanofil syringe (World Precision Instruments) mounted in a Quintessential Stereotaxic Injector (Stoelting) controlled by a Motorized Lab Standard Stereotax (Harvard Apparatus). Left or right hemisphere was balanced across all animals. One week later, the rat was again induced under anesthesia and 4 skull screws were implanted to provide stable hold for the subsequent implant. The viral craniotomy was reopened to a diameter of 1.8 mm, and cortical tissue and corpus callosal fibers above the hippocampus were aspirated away using a 27 and 30 gauge blunt needle. Following this aspiration, and assuring no bleeding persisted in the craniotomy, a stack of two 1.8 mm diameter Gradient Refractive INdex (GRIN) lenses (Edmund Optics) was implanted over the hippocampus and cemented in place with methacrylate bone cement (Simplex-P, Stryker Orthopaedics). The dorsal surface of the skull and bone screws were cemented with the GRIN lens to ensure stability of the implant, while the dorsal surface of the implanted lens was left exposed.

Two to three weeks later, rats were again placed under anesthesia in order to cement a 3D printed baseplate above the lens. First a second GRIN lens was optically glued (Norland Optical Adhesive 68, Edmund Optics) to the surface of the implanted lens and cured with UV light. The pitch of each GRIN lens was ≲0.25, so implanting two stacked lenses yielded a total pitch of ≲0.5. This half pitch provides translation of the image at the bottom surface of the lenses to the top while maintaining the focal point below the lens. This relay implant enables access to tissue deep below the skull surface. The MiniLFOV was placed securely in the baseplate and then mounted to the stereotax to visualize the calcium fluorescence and tissue. The baseplate was then cemented in place above the relay lenses at the proper focal plane and allowed to cure.

### Linear alternation behavior

After rats had been baseplated, they were placed on food restriction to reach a goal weight of 85% *ad lib* weight and then began behavioral training. Time between the beginning of the surgical procedures and the start of behavior training was typically 6-8 weeks. Rats earned 20 mg chocolate pellets by alternating between two rewarded ends of a linear track (250 cm) during 15 min recordings beginning 5 days after baseplating. After receiving a reward at one end, the next reward had to be earned at the other end by crossing the center of the track.

### Behavior tracking

A webcam mounted in the behavior room tracked a red LED located on the top of the Miniscope and this video was saved alongside the calcium imaging via the Miniscope DAQ software with synchronized frame timestamps. These behavior video files were initially processed by custom python code, where all the session videos were concatenated together into one tiff stack, downsampled to 15 frames per second, the median of the stack was subtracted from each image, and finally they were all rescaled to the original 8-bit range to yield the same maximum and minimum values before subtraction. Background subtracted behavior videos were then processed in MATLAB. The rat’s position in each frame was determined using the location of the red LED on the camera. Extracted positions were then rescaled to remove the camera distortion and convert the pixel position to centimeters according to the maze size. Positional information was then interpolated to the timestamps of the calcium imaging video using a custom MATLAB script.

### Histology

At the end of the experiment, rats were anesthetized with isoflurane, intraperitoneally injected with 1 mL of pentobarbital, then transcardially perfused with 100 mL of 0.01M PBS followed by 200 mL of 4% paraformaldehyde in 0.01M PBS to fix the brain tissue. Brains were sectioned at 40 μm thickness on a cryostat (Leica), mounted on gelatin prepared slides, then imaged on a confocal microscope (Zeiss) to confirm GFP expression and GRIN lens placement.

## Supplementary Video Captions

**Supplementary Video 1**. *Real-time motion correction*. The left and right windows show sensor video data before and after motion correction, respectively. The line graphs at bottom show x (yellow) and y (green) components of the image displacement between frames before (left) and after (right) motion correction.

**Supplementary Video 2**. *RTI view of contour-based calcium trace extraction*. The online image display (left window) shows the motion corrected and enhanced (i.e., background subtracted) sensor image data as it arrives in real time from the Ultra96 board. The right window shows a scrolling display of 63 selected calcium traces from regions outlined by colored borders in the image display window. These traces are derived on the Ultra96 by summing fluorescence within their respective contour regions, and the resulting trace values are transmitted (along with sensor image data) via ethernet to the host PC for display in the RTI window. For demonstration purposes, traces shown in the window on the right are normalized within the ranges of their own individual minimum and maximum values.

**Supplementary Video 3**. *Real-time decoding of contour-free calcium traces*. The online image display (left window) shows the motion corrected and enhanced mosaic of contour-free pixel mask tiles. The right window shows a scrolling heatmap display of 103 (out of the total 900) calcium traces with the highest tuning curve similarity scores, *S* (see main text). The calcium trace rows in the heatmap are sorted by the peak activity location of each trace, so that calcium activity can be seen to propagate through the population as the rat runs laps on the linear track. The line graph at the top shows the rat’s true position (blue line) together with its predicted position (orange line) decoded in real time by the linear classifier. For demonstration purposes, trace intensities shown in the scrolling heatmap are normalized within the ranges of their own individual minimum and maximum values.

## References

1. Ghosh, K. K. et al. Miniaturized integration of a fluorescence microscope. Nat. Methods 8, (2011).

2. Ziv, Y. et al. Long-term dynamics of CA1 hippocampal place codes. Nat. Neurosci. (2013) doi:10.1038/nn.3329.

3. Cai, D. J. et al. A shared neural ensemble links distinct contextual memories encoded close in time. Nature (2016) doi:10.1038/nature17955.

4. Aharoni, D., Khakh, B. S., Silva, A. J. & Golshani, P. All the light that we can see: a new era in miniaturized microscopy. Nature Methods (2019) doi:10.1038/s41592-018-0266-x.

5. Hart EE, Blair GJ, O’Dell TJ, Blair HT, Izquierdo A. Chemogenetic Modulation and Single-Photon Calcium Imaging in Anterior Cingulate Cortex Reveal a Mechanism for Effort-Based Decisions. (2020) J Neurosci. 40:5628–5643. doi: 10.1523/JNEUROSCI.2548-19.2020. PMID: 32527984; PMCID: PMC7363467.

6. Friedrich J, Zhou P, Paninski L. Fast online deconvolution of calcium imaging data. PLoS Comput Biol. 2017 Mar 14;13(3):e1005423. doi: 10.1371/journal.pcbi.1005423. PMID: 28291787; PMCID: PMC5370160.

7. Mitani A, Komiyama T. Real-Time Processing of Two-Photon Calcium Imaging Data Including Lateral Motion Artifact Correction. Front Neuroinform. 2018 Dec 18;12:98. doi: 10.3389/fninf.2018.00098. PMID: 30618703; PMCID: PMC6305597.

8. Chen Z, Blair HT, Cong J. (2019). LA-NorRMCorre: LSTM-Assisted Non-Rigid Motion Correction on FGPA for Calcium Image Stabilization. 27th ACM/SIGDA International Symposium on Field-Programmable Gate Arrays (FPGA), 1–6 (02/24/2019).

9. Chen Z, GJ Blair, Blair HT, Cong J. (2020). BLINK: bit-sparse LSTM inference kernel enabling efficient calcium trace extraction for neurofeedback devices. Proceedings of the ACM/IEEE International Symposium on Low Power Electronics. ISBN:978-1-4503-7053-0

10. Chen Z, Zhou J, Blair GJ, Blair HT, Cong J. “Efficient kernels for real-time position decoding from in vivo calcium images.” IEEE International Symposium on Circuits and Systems (ISCAS), (2022).

11. Friedrich J, Giovannucci A, Pnevmatikakis EA. Online analysis of microendoscopic 1-photon calcium imaging data streams. PLoS Comput Biol. 2021 Jan 28;17(1):e1008565. doi: 10.1371/journal.pcbi.1008565. PMID: 33507937; PMCID: PMC7842953.

12. Aharoni, D. & Hoogland, T. M. Circuit investigations with open-source miniaturized microscopes: Past, present and future. Frontiers in Cellular Neuroscience (2019) doi:10.3389/fncel.2019.00141.

13. Grosenick L, Marshel JH, Deisseroth K. Closed-loop and activity-guided optogenetic control. Neuron. 2015 Apr 8;86(1):106–39. doi: 10.1016/j.neuron.2015.03.034. PMID: 25856490; PMCID: PMC4775736.

14. Zhang Z, Russell LE, Packer AM, Gauld OM, Häusser M. Closed-loop all-optical interrogation of neural circuits in vivo. Nat Methods. 2018 Dec;15(12):1037–1040. doi: 10.1038/s41592-018-0183-z. Epub 2018 Nov 12. PMID: 30420686; PMCID: PMC6513754.

15. Benjamin B. Scott, Stephan Y. Thiberge, Caiying Guo, D. Gowanlock R. Tervo, Carlos D. Brody, Alla Y. Karpova, David W. Tank (2018). Imaging Cortical Dynamics in GCaMP Transgenic Rats with a Head-Mounted Widefield Macroscope. Neuron, 100:1045–58.

16. de Groot, A. et al. (2020). Ninscope, a versatile miniscope for multi-region circuit investigations. Elife doi:10.7554/eLife.49987.

17. Guo C, Blair G, Sehgal M, Jimka FNS, Bellafard A, Silva AJ, Golshani P, Basso MA, Blair HT, Aharoni D. Miniscope-LFOV: A large field of view, single cell resolution, miniature microscope for wired and wire-free imaging of neural dynamics in freely behaving animals (submitted).

18. O’Keefe, J., and J. Dostrovsky. 1971. “The Hippocampus as a Spatial Map. Preliminary Evidence from Unit Activity in the Freely-Moving Rat.” Brain Research 34 (1): 171–175. doi:10.1016/0006-8993(71)90358-1.

19. Wilson MA, McNaughton BL. Dynamics of the hippocampal ensemble code for space. Science. 1993 Aug 20;261(5124):1055-8. doi: 10.1126/science.8351520. Erratum in: Science 1994 Apr 1;264(5155):16. PMID: 8351520.

20. Giovannucci, Andrea, Johannes Friedrich, Pat Gunn, Jérémie Kalfon, Brandon L Brown, Sue Ann Koay, Jiannis Taxidis, et al. 2019. “CaImAn an Open Source Tool for Scalable Calcium Imaging Data Analysis.” ELife 8:e38173. doi:10.7554/elife.38173.

21. Deng X, Liu DF, Kay K, Frank LM, Eden UT (2015). Clusterless Decoding of Position from Multiunit Activity Using a Marked Point Process Filter. Neural Comput. 27(7):1438–1460. doi:10.1162/NECO_a_00744

22. Lu J, Li C, Singh-Alvarado J, Zhou ZC, Fröhlich F, Mooney R, Wang F. MIN1PIPE: A Miniscope 1-Photon-Based Calcium Imaging Signal Extraction Pipeline. Cell Rep. 2018 Jun 19;23(12):3673–3684. doi: 10.1016/j.celrep.2018.05.062. PMID: 29925007; PMCID: PMC6084484.

23. Dana, H., Sun, Y., Mohar, B. et al. High-performance calcium sensors for imaging activity in neuronal populations and microcompartments. Nat Methods 16, 649–657 (2019). https://doi.org/10.1038/s41592-019-0435-6

24. Buzsáki, G. (2015). Hippocampal sharp wave-ripple: A cognitive biomarker for episodic memory and planning. Hippocampus, 25(10), 1073–1188. https://doi.org/10.1002/hipo.22488

25. Liu C, Li M, Wang R, Cui X, Jung H, Halin K, You H, Yang X, Chen W. Online Decoding System with Calcium Image From Mice Primary Motor Cortex. Annu Int Conf IEEE Eng Med Biol Soc. 2021 Nov;2021:6402–6405. doi: 10.1109/EMBC46164.2021.9630138. PMID: 34892577.

26. M. Taniguchi et al., “Open-Source Software for Real-time Calcium Imaging and Synchronized Neuron Firing Detection,” 2021 43rd Annual International Conference of the IEEE Engineering in Medicine & Biology Society (EMBC), 2021, pp. 2997–3003, doi: 10.1109/EMBC46164.2021.9629611.

27. Tu, M., Zhao, R., Adler, A., Gan, W. B., & Chen, Z. S. (2020). Efficient Position Decoding Methods Based on Fluorescence Calcium Imaging in the Mouse Hippocampus. Neural computation, 32(6), 1144–1167. https://doi.org/10.1162/neco_a_01281

28. Kinsky NR, Sullivan DW, Mau W, Hasselmo ME, Eichenbaum HB. Hippocampal Place Fields Maintain a Coherent and Flexible Map across Long Timescales. Curr Biol. 2018 Nov 19;28(22):3578-3588.e6. doi: 10.1016/j.cub.2018.09.037. Epub 2018 Nov 1. PMID: 30393037; PMCID: PMC6331214.

29. Pnevmatikakis EA, Soudry D, Gao Y, Machado TA, Merel J, Pfau D, Reardon T, Mu Y, Lacefield C, Yang W, Ahrens M, Bruno R, Jessell ™, Peterka DS, Yuste R, Paninski L (2016) Simultaneous denoising, Deconvolution, and demixing of calcium imaging data Neuron 89:285–299. https://doi.org/10.1016/j.neuron.2015.11.037

